# Loss of 7-Dehydrocholesterol Reductase-mediated cholesterol biosynthesis activates IRF3 and inhibits control of *Mycobacterium marinum* infection

**DOI:** 10.64898/2026.05.07.723652

**Authors:** Xiangyu Sui, Darryl JY Han, David M Costa, Vinitha A Jacob, Stefan H Oehlers

## Abstract

Cholesterol immunometabolism is a critical controller of immunopathology in respiratory infections such as tuberculosis. Smith-Lemli-Opitz syndrome (SLOS) patients are affected by a loss of 7-dehydrocholesterol reductase (DHCR7) function and have elevated 7-dehydrocholesterol (7DHC) and reduced cholesterol. Increased 7DHC has been found to be protective against viral infections in a range of infection models however SLOS patients have a higher susceptibility to respiratory infection. Here we use the zebrafish-*Mycobacterium marinum* infection model to demonstrate a compromised innate immune response to bacterial infection in the absence of *dhcr7*. We correlate increased 7DHC with increased activation of the IRF3/type I interferon axis and demonstrate Irf3 is a targetable signaling node to restore anti-bacterial immunity in a *dhcr7*-depleted background.

**Plain English summary:** Loss of 7-dehydrocholesterol reductase causes Smith-Lemli-Opitz syndrome. One of the metabolic features of Smith-Lemli-Opitz syndrome is increased 7-dehydrocholesterol (7DHC). We find increased 7DHC inhibits the ability of zebrafish to control mycobacterial infection by mis-activating an antiviral immune response at the expense of a protective anti-bacterial immune response. Our study suggests the susceptibility to respiratory infections and increased neuroinflammation in Smith-Lemli-Opitz syndrome could be treated by targeting the antiviral protein IRF3.

## Introduction

7-dehydrocholesterol reductase (DHCR7) is one of the key enzymes involved in the cholesterol synthesis pathway, converting 7-dehydrocholesterol (7DHC) to cholesterol. Loss of function in *DHCR7* causes Smith–Lemli–Opitz syndrome (1), an inborn error of cholesterol synthesis with heightened susceptibility to respiratory infections. Although infection susceptibility is primarily thought to be the result of structural and reduced tonicity in the respiratory tract, there is building evidence for innate immune defects in the absence of DHCR7 with recent mouse model papers demonstrating pulmonary neutrophil recruitment defects, thought to be mediated by reduced TLR signalling, and excessive immune activation in astrocytes and microglia (2, 3). Conversely, viral infection studies revealed that DHCR7 inhibition-induced 7DHC accumulation activates Interferon regulatory factor 3 (IRF3), driving type I interferon expression and enhancing antiviral defences (4).

Host lipid metabolism is a complex process modulated by host immune and pathogen processes. In the context of tuberculosis (TB), caused by infection with *Mycobacterium tuberculosis*, the accumulation of lipid species in macrophages has been demonstrated to both aid and hinder the immune response (5, 6). Furthermore, Type I interferon signalling has been found to be protective and detrimental across different TB model systems demonstrating the ambiguity of these DHCR7-related pathways in the context of infection (7).

The zebrafish-*Mycobacterium marinum* infection model has been extensively used to study TB, including alterations in host metabolism (8, 9). We have previously found infection-induced foam cell formation and dyslipidaemia are conserved in the zebrafish-*M. marinum* model (10, 11). Depletion of the low-density lipoprotein receptor or preventing thrombocyte-macrophage interactions reduced foam cell formation correlated with decreased *M. marinum* burden in zebrafish (11, 12). Here we sought to investigate the Dhcr7-cholesterol metabolism-Irf3/type I interferon axis in a zebrafish model of acute mycobacterial infection.

## Methods

### Zebrafish husbandry and infection procedure

Zebrafish embryos were obtained by natural spawning of adult zebrafish and maintained at 28 °C in E3 embryo media supplemented with methylene blue. Embryos were dechorionated using pronase and transferred to E3 medium containing 45 µg/mL 1-phenyl-2-thiourea (Sigma-Aldrich) from one day post-fertilization (dpf). *M. marinum* was cultured and prepared for infection as previously described (13). At 2 dpf, embryos were anesthetized with 0.4% (w/v) tricaine prior to caudal vein microinjection of approximately 100 colony-forming units (CFU) of *M. marinum* for systemic infections. Following infection, embryos were recovered into PTU-supplemented E3 medium and maintained at 28 °C for 3-5 days post infection (dpi).

### CRISPR-Cas9 editing of *dhcr7* and *irf3*

Guide RNAs (gRNAs) were generated by *in vitro* transcription using a common scaffold primer 5′-AAAAGCACCGACTCGGTGCCACTTTTTCAAGTTGATAACGGACTAGCCTTATTTTAACTTGCTATTTCTA GCTCTAAAAC-3′ annealed to gene specific primers: *dhcr7* 5′-TAATACGACTCACTATAGGAAGGAGCCCGGTTCCAGAGTTTTAGAGCTAGAAATAGC-3′ 5′-TAATACGACTCACTATAGGAACTCCTCCTACGTAGCCGTTTTAGAGCTAGAAATAGC-3′ 5′-TAATACGACTCACTATAGGTGACAAACTCTATGATCCGTTTTAGAGCTAGAAATAGC-3′ 5′-TAATACGACTCACTATAGGATGAGGACCCCTGCTGCGGTTTTAGAGCTAGAAATAGC-3′, *irf3* 5′-TAATACGACTCACTATAGGCAGTGCTCTTCGCGCTAAGTTTTAGAGCTAGAAATAGC-3′ 5′-TAATACGACTCACTATAGGAAGGTGCTGTCGGTGGTTGTTTTAGAGCTAGAAATAGC-3′ 5′-TAATACGACTCACTATAGGGTTGGTGGAGAACGACTCGTTTTAGAGCTAGAAATAGC-3′ 5′-TAATACGACTCACTATAGGATTCACGGTCACCGTTGGGTTTTAGAGCTAGAAATAGC-3′ as previously described (14).

### High resolution melt curve analysis

Editing efficiency of gRNAs targeting *dhcr7* was verified using MeltDoctor^™^ HRM Master Mix (Thermofisher), and an amplicon generated by forward primer 5′-CTTTCGGGTGTGATCCTGCT -3′ and reverse primer 5′-AAATGTGACCCAGATGGCGT -3′.

### Lipid mass spectroscopy

Pools of 12 embryos were homogenised in ice cold methanol by bead beating, a 1:2 ratio of chloroform was added to the methanol to solubilise lipids and samples were stored at -80 °C prior to analysis. A 7-dehydrodesmosterol (7DHC) standard (Toronto Research Chemicals) was added to an in-house cholesterol mass spectroscopy panel at the Duke-NUS Metabolomics Core for detection of cholesterol and 7DHC.

### Fluorescence imaging and measurement

Embryos were anesthetized with tricaine, mounted in methylcellulose gel for imaging on either a Nikon SMZ25 stereoscope or an inverted Nikon Eclipse Ti2, and images were analysed by fluorescent pixel count as previously described (13).

### cGAMP injection

3’3’-cyclic GAMP (cGAMP) Fluorinated (InvivoGen) was dissolved in water at a concentration of 50 mg/ml. 1 nl was injected into the circulation of 2 dpf zebrafish embryos as per the infection procedure. cGAMP-injected embryos were recovered into PTU-supplemented E3 medium and maintained at 28 °C for 3 days prior to imaging.

### Gene expression

RNA was extracted from pools of 15-30 infected embryos using Trizol (Thermofisher) and cDNA synthesis was carried out with the Applied Biosystems High Capacity cDNA synthesis kit (Thermofisher). Gene expression was normalized to the zebrafish *18S* rRNA gene using the primer set 5′-TCGCTAGTTGGCATCGTTTATG-3′ and 5′-CGGAGGTTCGAAGACGATCA-3′. Primer sets for genes of interest: irf3 5′-ACTGCCATTCCTGAGCCAAA-3′ and 5′-TCGTTCTCCACCAACTGCTC-3′, *ifnphi1* 5′-CTACTTGCGAATGGCTTGGC-3′ and 5′-CCTGGTCCTCCACCTTTGAC-3′, *ifnphi3* 5′-GAGAGGCTTCGCTCAAGGTT-3′ and 5′-GGGTGCGGAGAATCTCACTT-3′.

### Statistical analysis

All statistical analysis were performed using GraphPad Prism. T tests and one-way ANOVA analysis were carried out for pairwise comparisons and comparisons of three or more groups, respectively.

## Results

To investigate the role of Dhcr7 in immunity to mycobacterial infection, *dhcr7* was knocked down in zebrafish embryos using CRISPR/Cas9 mutagenesis and embryos were infected with fluorescent *M. marinum* (Figure 1A). Editing efficiency was verified by high resolution melt curve analysis (Figure 1B). As expected from the conserved role of DHCR7 in cholesterol synthesis, knockdown of *dhcr7* significantly reduced cholesterol production and increased the relative abundance of the 7DHC precursor metabolite (Figure 1C). Analysis of infection burden revealed increased *M. marinum* infection burden in *dhcr7* crispants (Figure 1D), suggesting *dhcr7* plays a protective role during *M. marinum* infection.

**Figure 1:**
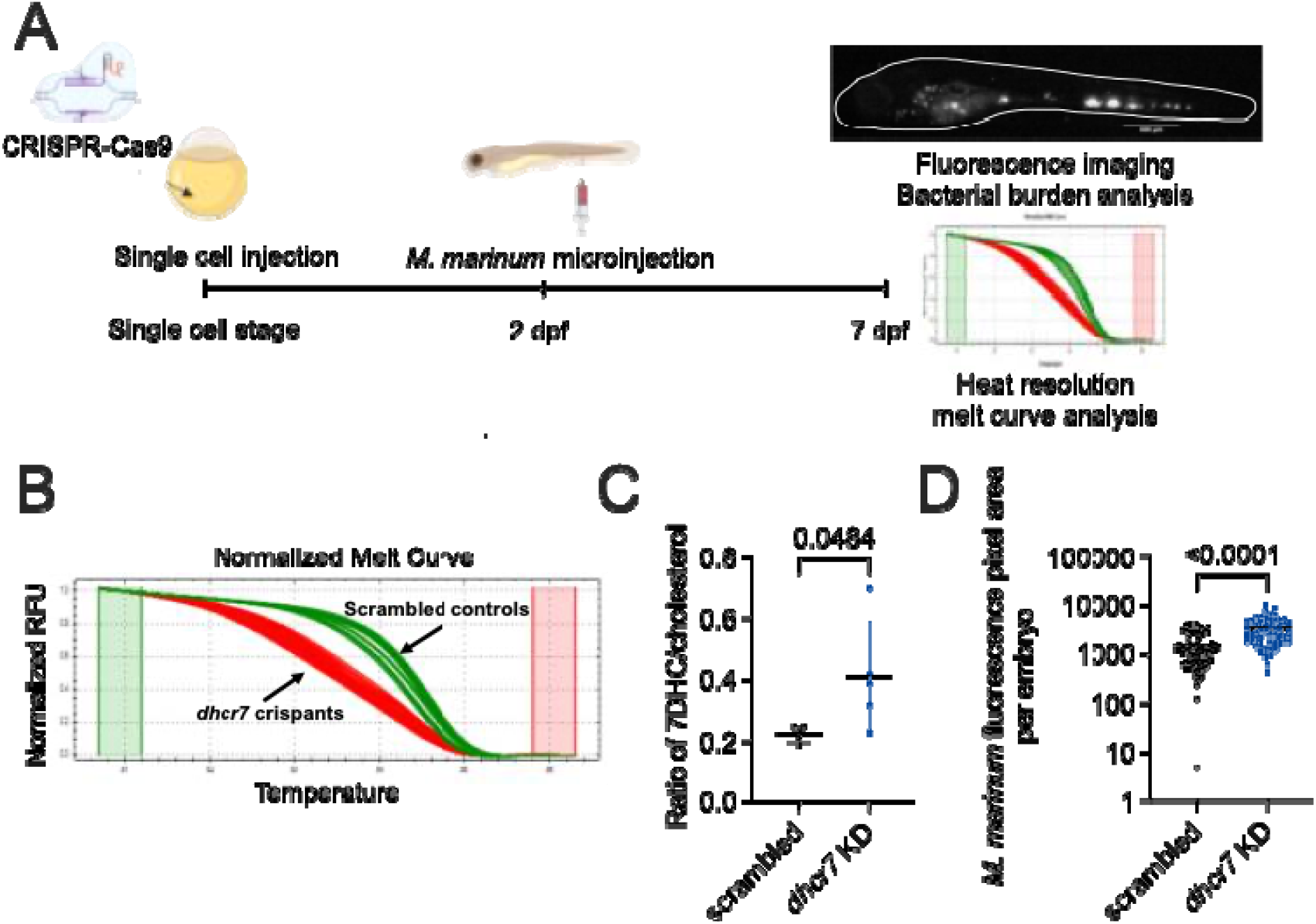
Knockdown of *dhcr7* increases 7DHC and *M. marinum* infection burden. A. Workflow for zebrafish embryo experiments generating *dhcr7* crispant embryos and analysis of *M. marinum* infection burden by imaging. B. Example of HRM determination of gene editing in *dhcr7* crispant embryos. C. Quantification of 7DHC and cholesterol from 5 dpi *dhcr7* crispant embryos. Each dot represents one pool of embryos from biological replicates. D. Quantification of *M. marinum* burden in 3 dpi *dhcr7* crispant embryos. Each dot represents a single animal, data is representative of three biological replicates.

To investigate whether 7DHC accumulation activates the IRF3/type I interferon signalling axis in our *M. marinum* infection system, we first visualized type I IFN pathway activity using the transgenic zebrafish reporter line *Tg(isg15:EGFP,myl7:EGFP)*^*z207*^, in which *isg15* promoter activity serves as a downstream readout of type I IFN activation (15). As *M. marinum* infection did not produce detectable *isg15:GFP* fluorescence (Figure 2A), cGAMP was injected into embryos to stimulate type I IFN signaling through the STING pathway. In the cGAMP-injected groups, *isg15* promoter activity was detected in the distal kidney region at 3 days post-injection (Figure 2B). Quantification of the fluorescent signal in the distal kidneys of *Tg(isg15:EGFP,myl7:EGFP)*^*z207*^ embryos revealed that *dhcr7* knockdown increased the area of cGAMP-induced *isg15* promoter driven GFP fluorescence, compared to scrambled controls, consistent with the previously published role of 7DHC accumulation in driving IRF3 activation (Figure 2B).

**Figure 2:**
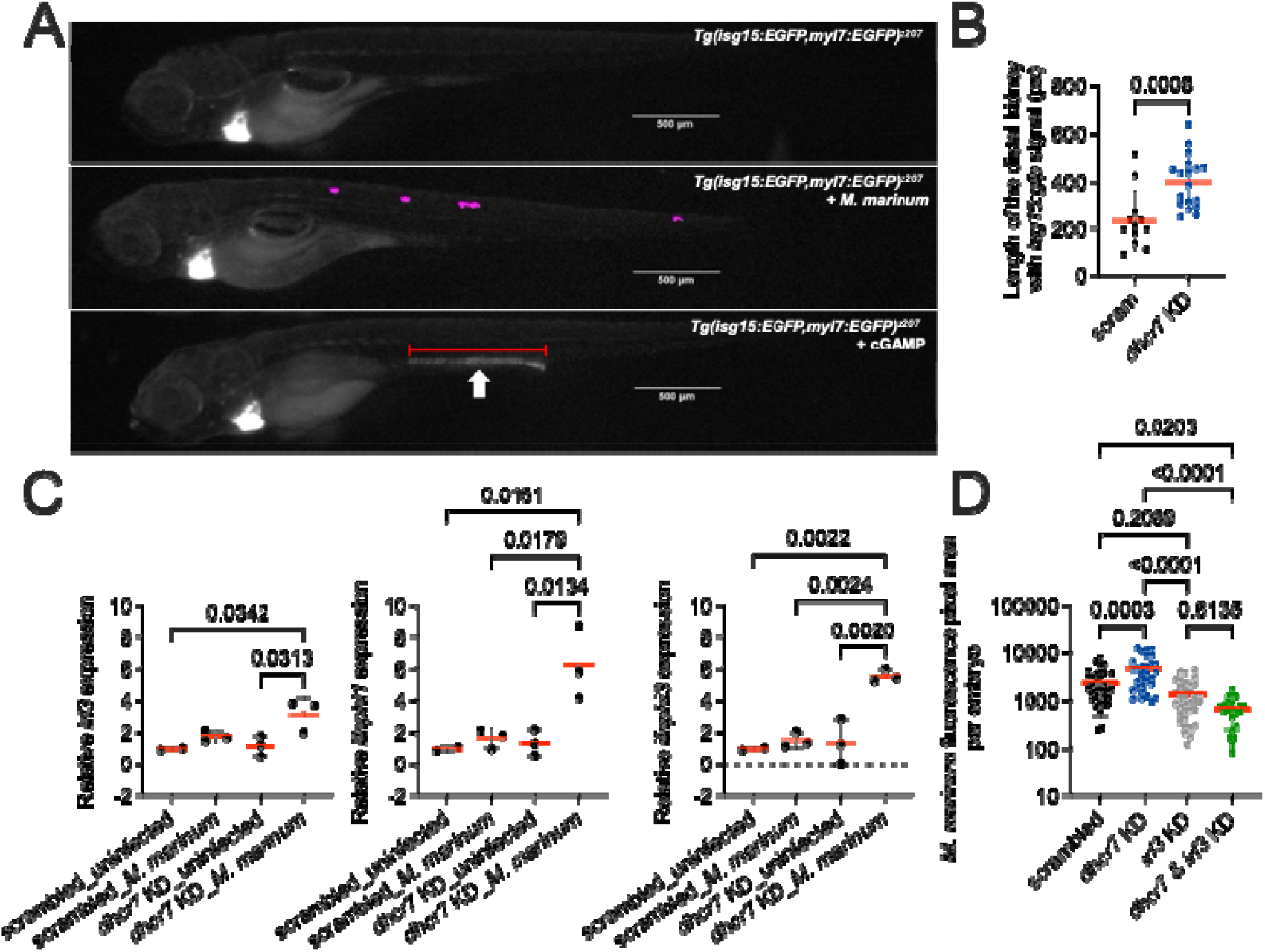
*dhcr7* depletion hyperactivates the IRF3/type I IFN signaling axis during *M. marinum* infection. A. Representative images of 3 day post injection *Tg(isg15:EGFP,myl7:EGFP)*^*z207*^ zebrafish embryos infected with *M. marinum* (magenta) or injected with cGAMP. Scale bar represents 500 µm. Arrow indicates location of cGAMP-responsive distal kidney. B. Quantification of cGAMP-induced *isg15:gfp* fluorescent area in the distal kidney of 3 day post injection *dhcr7* crispant embryos. Each dot represents a single embryo, data is representative of three biological replicates. C. Quantification of *irf3, ifnphi1*, and *ifnphi3* transcripts by qPCR in 5 dpi *dhcr7* crispant embryos. Each dot represents one pool of embryos from biological replicates. D. Quantification of *M. marinum* burden in 5 dpi *dhcr7* and *irf3* double crispant embryos. Each dot represents a single embryo, data is representative of three biological replicates.

To more sensitively assess type I IFN activation, canonically downstream of Irf3, we next quantified the transcription of *irf3* and type I IFN genes by qPCR in *M. marinum*-infected *dhcr7* crispant embryos. In zebrafish, type I interferons are referred to as *ifnphi*, with *ifnphi1* and *ifnphi3* being the main isoforms expressed during the embryonic stage (16). qPCR results showed that irf3, ifnphi1, and ifnphi3 were significantly upregulated in *dhcr7* knockdown embryos infected with *M. marinum*, compared to uninfected and infected scrambled controls (Figure 2C).

Next, *irf3* knockdown and *dhcr7/irf3* double knockdown were performed to further validate the involvement of IRF3 in susceptibility to *M. marinum* infection. Bacterial burden analysis showed that the increased susceptibility to *M. marinum* infection observed in the *dhcr7* knockdown group was reversed in the double knockdown group (Figure 2D), consistent with *dhcr7* depletion-induced 7DHC accumulation driving susceptibility to *M. marinum* infection by hyperactivation of the IRF3/type I IFN signalling axis.

## Discussion

Our study reveals a previously uncharacterised role of *dhcr7* in suppressing detrimental Irf3-driven responses during mycobacterial infection. These data suggest inappropriate IRF3 activation may contribute to the increased respiratory bacterial infections by experienced by SLOS patients. IRF3 primarily mediates anti-viral immunity and the quality of inflammation mediated by chronic IRF3 activation drives immunopathology to a range of important respiratory pathogen including *M. tuberculosis* and SARS-CoV-2 (7, 17), thus IRF3 activation may represent a therapeutic target in the management of SLOS susceptibility to respiratory infections and neuroinflammation.

Our findings contribute to understanding the complex role of cholesterol synthesis intermediates in modulating host immunity. The zebrafish is increasingly proving to be an excellent *in vivo* model of human cholesterol-infection biology exemplified by a recent study contrasting the previously reported antiviral role of the cholesterol 25-hydroxylase metabolite 25-hydroxycholesterol with exacerbated brain endothelial dysfunction in a model of SARS-CoV-2 infection (18, 19). Together with our data linking DHCR7 to septic shock (20), these studies demonstrate the power of *in toto* analysis of cholesterol metabolism in immunopathogenesis.

## List of abbreviations

7DHC: 7-dehydrocholesterol
DHCR7: 7-dehydrocholesterol reductase
dpf: Days post fertilisation
dpi: Days post infection
gRNA: Guide RNA
IRF3: Interferon regulatory factor 3
SLOS: Smith–Lemli–Opitz syndrome
TB: Tuberculosis

## Declarations

### Ethics approval

Animal experiments were approved by the A*STAR Institutional Animal Care and Use Committee under protocols 211667 and 221694.

### Consent for publication

Not applicable.

### Availability of data and materials

Primary research data is available upon reasonable request to the corresponding author.

### Author’s contributions

VAJ and SHO conceived the project, XS and DJYH designed methodology, XS and DMC performed experiments, SHO wrote the manuscript, all authors edited and approved the manuscript.

## Acknowledgements

The authors thank the A*STAR IMCB Aquarium Platform for expert zebrafish husbandry, Sakshi Agarwal and Amit Singhal for discussion on cholesterol and TB, Dexter Xiao and Chuan Yan for discussion on zebrafish interferon signalling, and Kaiwen Chen and Yongliang Zhang for supervision.

